# Hormonal Modulation of Subjective Sleep Quality and Perivascular Space Volume Across the Menstrual Cycle: An Observational Dense-Sampling Study

**DOI:** 10.64898/2026.09.04.749189

**Authors:** Daryna Apter, Jorik D. Elberse, Laura Stankeviciute, Susanne Weis, David Elmenhorst, Jeffrey Iliff, Simon B. Eickhoff, Carina Heller, Masoud Tahmasian

## Abstract

**Background:** Hormonal fluctuations during the menstrual cycle (and their suppression) are accompanied by altered mood, cognition, and sleep, yet their relationship to subjective sleep quality and glymphatic function - a sleep-dependent brain waste-clearance pathway that involves perivascular spaces (PVS) - is unclear.

**Methods:** We examined daily oestradiol, progesterone, luteinizing hormone (LH), follicle-stimulating hormone (FSH), subjective sleep quality, and MRI-derived PVS volume (PVSV) in a single healthy woman across two 30-day periods: during her physiological menstrual cycle and, one year later, under oral contraceptive pill (OCP) use. We employed linear regression models to examine whether hormones are associated with sleep quality and PVSV, and whether they moderate the sleep quality–PVSV association during the physiological menstrual cycle. Subsequently, we conducted permutation-based Spearman rank correlations to assess whether day-to-day changes in sleep quality were associated with corresponding changes in PVSV across the physiological menstrual cycle and OCP use. False discovery rate (FDR) correction was used to control for multiple comparisons.

**Findings:** Higher LH levels were associated with poorer sleep quality during the physiological cycle (b=2·15, 95% CI 0·64 to 3·67, p_FDR_=0·028), while FSH moderated the sleep quality–PVSV association (b=−92·98, 95% CI −160·15 to −25·81, p_FDR_=0·034), suggesting that lower FSH was associated with lower PVSV, whereas this association reversed at elevated FSH levels characteristic of the periovulatory window. During the physiological menstrual cycle, day-to-day decreases in sleep quality were associated with synchronous increases in PVSV, driven by centrum semiovale PVSV (ρ=0·55, 95% CI 0·30 to 0·67, p_perm_=0·0095). This association was absent during OCP use and differed between hormonal milieus (Δρ=0·72, p=0·040).

**Interpretation:** Ovulation is characterized by poor sleep quality and reduced PVSV, potentially reflecting noradrenergic modulation. The association between sleep quality and PVSV day-to-day changes is shaped by the hormonal milieu, being absent under OCP. PVSV increase associated with poor sleep may reflect elevated sleep pressure promoting vasomotor-mediated cerebrospinal fluid inflow.

**Funding:** None

## Introduction

The menstrual cycle is characterised by rhythmic fluctuations in gonadotropic and ovarian hormones that coordinate ovulation and endometrial remodelling. These fluctuations are accompanied by alterations in mood, cognition, and sleep patterns.^1^ Sleep quality is particularly reduced during the menstrual, mid-cycle, and late luteal phases,^2^ with hormonal fluctuations thought to orchestrate these changes. Elevated progesterone during the luteal phase promotes increased daytime sleepiness, an effect commonly attributed to the GABAergic neurosteroid allopregnanolone.^3^ Conversely, the periovulatory surge in gonadotropins has been associated with more nocturnal awakenings.^3^ Pharmacological suppression of hormonal fluctuations, e.g., via oral contraceptive pills (OCPs), may stabilise sleep quality by reducing endocrine variability, though findings remain divergent, likely attributable to heterogeneity in OCP formulations.^4^

Sleep health is a fundamental pillar of brain health, and poor sleep quality is associated with a range of neurological and psychiatric conditions.^5^ The glymphatic system (GS) has been proposed as a potential modulator of sleep-related brain health, comprising a brain waste-clearance system that is active during non-rapid eye movement (NREM) sleep.^6^ It is proposed to constitute a perivascular network that promotes cerebrospinal fluid (CSF) flow along the perivascular spaces (PVS) of penetrating arteries, enabling CSF– interstitial fluid exchange and clearance of metabolic by-products, including amyloid-β.^7^ Noradrenergic signalling from the locus coeruleus (LC), a brainstem nucleus regulating wakefulness and sleep, was shown to modulate CSF flow through rhythmic norepinephrine release during NREM sleep, which drives slow vasomotion that pumps CSF from the PVS into the brain parenchyma.^8^ Glymphatic activity can be measured invasively by intrathecal injection of contrast agents and tracking their distribution in the brain via magnetic resonance imaging (MRI).^9^ Given the invasive nature and limited feasibility of such approaches in retrospective study designs, PVS volume (PVSV) gained attention as a putative correlate of CSF inflow.^10^ Due to its vascular proximity, PVSV is shaped by alterations in arterial inflow pathways,^10^ possibly linking reduced PVSV to increased resistance to CSF inflow.^11^ Notably, ovarian hormones are known to modulate cerebrovascular properties, as reflected in cyclical changes in cerebrovascular reactivity across the menstrual cycle.^12^ Conversely, OCP-induced suppression of reproductive hormone fluctuations has been linked to increased arterial stiffness,^13^ which may attenuate arterial pulsatility and thereby compromise perivascular flow, suggesting that both endogenous hormonal fluctuations and their pharmacological suppression may alter perivascular fluid dynamics and PVSV.

Collectively, existing evidence suggests that fluctuations in female reproductive hormones may affect sleep quality, and sleep represents a key determinant of glymphatic function. However, whether endogenous hormonal fluctuations and their suppression are associated with concurrent changes in subjective sleep quality and PVSV remains unknown. Addressing this question requires dense longitudinal within-person designs to disentangle sustained endocrine states from day-to-day dynamic hormonal variation. We therefore leveraged dense daily sampling of circulating reproductive hormones, subjective sleep quality assessments, and MRI acquisitions of one subject across 30 consecutive days spanning both a physiological menstrual cycle and OCP use (**Figure 1**). We aimed to determine whether endogenous hormonal fluctuations during the physiological menstrual cycle (*i*) are associated with subjective sleep quality and PVSV; (*ii*) whether they moderate the association between sleep quality and PVSV. We further explored (*iii*) the role of hormonal suppression (OCP use) on the observed patterns.

**Figure 1:**
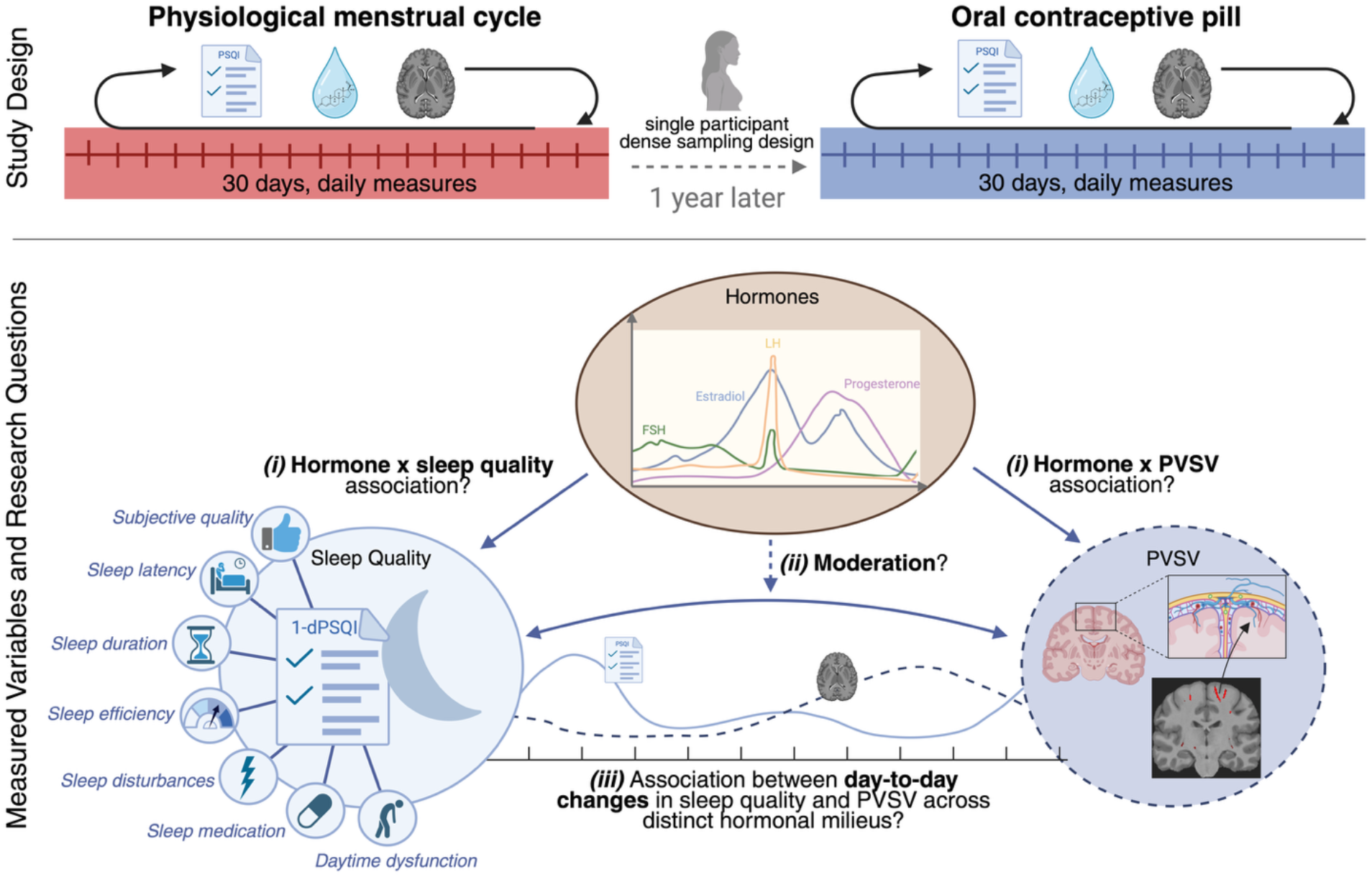
Study design and research questions. **Upper panel:** A healthy young woman underwent daily blood sampling, subjective sleep quality assessments (1-dPSQI), and structural MRI on 30 consecutive days during a physiological menstrual cycle and one year later while taking an oral contraceptive pill (initiated 10 months before the onset of the testing period). PVSV was derived from structural MRI scans. **Lower panel:** Three research questions were addressed: whether endogenous hormonal fluctuations during the physiological menstrual cycle (*i*) are associated with subjective sleep quality and PVSV; (*ii*) moderate the association between subjective sleep quality and PVSV; and (*iii*) the day-to-day variability in subjective sleep quality and PVSV, as well as the association between both measures, differs between two distinct hormonal milieus. 1-dPSQI, daily-adapted Pittsburgh Sleep Quality Index; PVSV, perivascular space volume. This figure was created with BioRender (BioRender.com/5o6vp7h).

## Methods

### Study Design & Data

This study utilised the publicly available, densely sampled longitudinal “28andMe” dataset available via the OpenNeuro repository (Accession No. ds002674, version 1.0.6; https://doi.org/10.18112/openneuro.ds002674.v1.0.6). The original study design and data acquisition procedures have been described in detail elsewhere.^14^ A brief outline of the experimental design is summarised below.

A 23-year-old Caucasian female underwent daily testing across two 30-day cycles. The participant had no history of neuropsychiatric or endocrine disorders or traumatic brain injury. She had regular menstrual cycles, with no missed periods and a stable cycle duration of 26–28 days. Prior to the first 30-day testing phase, hormone-based medication intake was excluded. The Human Subjects Committee of the University of California, Santa Barbara approved this design. The first testing day was scheduled independently of the cycle phase, which was subsequently determined based on hormonal assays. Before each session, the participant abstained from caffeinated beverages and maintained a 2-hour fast and fluid restriction. Testing began with daily questionnaires (9:00), followed by a blood draw (10:00 ± 30 minutes) and a subsequent one-hour MRI session. One year later, the 30-day testing protocol was repeated under identical conditions, this time during OCP use (0.02 mg ethinyl oestradiol, 0.1 mg levonorgestrel; Aubra, Afaxys Pharmaceuticals), initiated 10 months before the onset of the testing period.

### Hormonal Measures

Daily venous blood draws were performed by a licensed phlebotomist to determine the serum levels of gonadal hormones (oestradiol, progesterone) as well as the pituitary gonadotropins luteinising hormone (LH) and follicle-stimulating hormone (FSH). Steroid hormones were determined using liquid chromatography–tandem mass spectrometry and gonadotropins using an immunoassay at the Research Assay Core of Brigham and Women’s Hospital. Detailed methods for the blood analysis have been described elsewhere.^14^

### Subjective Sleep Quality Assessment

Subjective sleep quality was assessed each morning following the night to be evaluated via a daily-adapted version of the Pittsburgh Sleep Quality Index (1-dPSQI)^15^, which encompasses seven domains: subjective sleep quality, sleep latency, sleep duration, habitual sleep efficiency, sleep disturbances, use of sleep medication, daytime dysfunction, and the global score resulting from the sum of all components. The global score ranges from 0 to 21, with higher scores reflecting poorer sleep quality.

### Neuroimaging Acquisition and Preprocessing

Neuroimaging acquisition protocols have been described elsewhere.^14^ Daily high-resolution anatomical MRI scans were acquired on a Siemens 3T Prisma using a 64-channel head coil and a T1-weighted MPRAGE sequence (TR=2500 ms, TE=2.31 ms, TI=934 ms, flip angle=7°, slice thickness=0.8 mm).

All neuroimaging preprocessing and analysis pipelines were executed within Singularity containers using Bash scripts. Preprocessing of the T1w scans was performed with fMRIPrep (version 25.1.3), a pipeline of the NiPreps suite, which comprises a collection of open-source software tools for the standardised and automated preprocessing of neuroimaging data. Specifically, fMRIPrep includes (1) bias field correction of the T1w images to reduce signal inhomogeneities; (2) brain extraction by removing non-brain tissue; (3) segmentation into grey matter (GM), white matter (WM), and cerebrospinal fluid (CSF); (4) transformation of the images into the MNI standard space via spatial normalisation; and (5) reconstruction of cortical surfaces with subsequent anatomical parcellation. To preserve session-specific variation in PVSV and prevent potential attenuation of longitudinal anatomical changes resulting from fMRIPrep’s default generation of an intra-subject reference image by combining T1w scans across sessions, a BIDS filter file was applied to enforce session-wise preprocessing and generate independent preprocessed outputs for each session. All anatomical scans and PVS segmentation masks were subjected to manual visual quality control to exclude artefacts or erroneous segmentations, and they met the quality criteria.

### PVSV Quantification

Detection of PVS in T1w images was performed via a containerised version of the ShiVAi pipeline. It is based on the SHIVA-PVS model, a U-Net-based network architecture trained on three-dimensional spatial features of PVS.^16^ The pipeline enables automated PVS segmentation without introducing rater bias. The model weights used for this purpose are openly available via the ShiVAi GitHub repository (https://github.com/pboutinaud/SHiVAi). The Desikan–Killiany parcellation from FreeSurfer was used to identify PVS clusters within the centrum semiovale (CSO), basal ganglia (BG), and hippocampus, regions where PVS are most commonly observed on MRI. Total PVSV was defined as the cumulative volume of all voxels labelled as PVS by the model within each anatomical region.

### Statistical Analysis

#### Data Preprocessing and Correction for Serial Dependence

All statistical analyses and data visualisation were performed using R (version 4.5.3). Before analysis, all hormonal variables were log-transformed to address their naturally right-skewed distribution. Given the limited sample size, outliers were identified using a predefined criterion (|z|>3 for PVSV or 1-dPSQI) to avoid disproportionate influence of single values on subsequent analyses, resulting in a single outlier (PSQI = 8, OCP condition). Where serial dependence violated the independence assumptions of standard parametric methods, we applied resampling-based inference, using moving-block permutation tests for significance assessment and moving-block bootstrap procedures for 95% confidence interval estimation. Both approaches resampled contiguous blocks of observations, thereby preserving the temporal structure of the data. Block length was selected according to the theoretical optimum for block resampling procedures at the given sample size (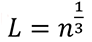; *L* = block length).^17^ Mean levels of all variables were compared between conditions using moving-block permutation (10,000 iterations, block length = 4 days). We conducted three sets of analyses, as follows:

#### 1) Hormone regression models for sleep quality and PVSV

Linear regression models were fitted exclusively in the physiological menstrual cycle condition to examine whether hormone levels predicted 1-dPSQI total scores and PVSV (PSQI ∼ Hormone; PVSV ∼ Hormone). Given that OCP alters endogenous hormonal profiles through exogenous hormone exposure, analyses were restricted to the physiological cycle to preserve the interpretability of endogenous hormone–outcome associations. To mitigate multicollinearity among hormonal trajectories, four separate models were fitted per outcome, each including a single hormone (oestradiol, progesterone, LH, or FSH). To further characterise hormone–sleep quality associations, exploratory post hoc Spearman rank correlations were computed between hormone levels and 1-dPSQI component scores. More details are in the Supplementary Materials.

#### 2) Hormone × sleep interaction models

To examine whether hormone levels moderated the sleep quality–PVSV association, interaction models were fitted for each hormone, with PVSV as the dependent variable (PVSV ∼ 1-dPSQI × Hormone). 1- dPSQI and hormone levels were mean-centred to reduce collinearity between main and interaction effects.

Regression models were evaluated for residual autocorrelation using the Durbin–Watson test. As residual autocorrelation was not significant, standard parametric inference was retained, and ordinary least squares (OLS) regression models were fitted. Residual normality was evaluated using the Shapiro–Wilk test and visual inspection of Q–Q plots. Cook’s distance was computed to identify influential observations, and models were re-evaluated after exclusion of identified influential observations, if present. To control the false discovery rate (FDR) arising from multiple comparisons, p-values were adjusted using the Benjamini–Hochberg procedure (FDR-adjusted p<0.05), applied separately within each model group. For all regression models, regression coefficients (b), standard errors (SE), and 95% confidence intervals (95% CI) are reported.

#### 3) Association of day-to-day changes between sleep quality and PVSV

To capture the amplitude of day-to-day variability in sleep quality and PVSV as well as the association between both measures across two distinct hormonal milieus, we employed day-to-day changes (Δ) in dPSQI and PVSV rather than their absolute levels. The association was assessed using Spearman rank correlations at the difference level (ΔPVSV and ΔPSQI), calculated separately for the physiological menstrual cycle and the OCP condition. Day-to-day changes were calculated as first-order differences between successive study days (*ΔX_t_* = *X_t_* − *X_t_*_−1_; where *t* represents the current measurement day). Given one identified outlier on 1-dPSQI in the OCP condition, two Δ values were excluded, corresponding to the transitions into and out of the affected day. Although autocorrelation was present in both delta series, AR(1) regression was deliberately avoided because modelling the autoregressive structure would partially absorb the short-term covariance of interest and thereby attenuate the association under investigation.^18^ Thus, moving-block permutation (block length = 3 days) was employed to permute Δ1-dPSQI while holding ΔPVSV constant, thereby preserving short-term serial dependence and generating an empirical null distribution for the association. Association strength was then compared between conditions using a block-permutation (block-length = 4 days) test of the difference in Spearman coefficients. To determine whether the Δ1-dPSQI–ΔPVSV association was specific to synchronous changes or extended to delayed effects, lagged cross-correlations were computed for lags of +1 to +3 days, based on the directional assumption that sleep quality influences PVSV.

To examine the contribution of each PVSV subregion to the overall Δ1-dPSQI–ΔPVSV association, region-specific analyses (CSO and BG) were performed for any condition exhibiting a significant Δ1-dPSQI– ΔPVSV association, following the procedure described above. Unique regional contributions were then quantified using partial Spearman correlations while controlling for the complementary PVSV subregion, with statistical significance assessed by block-permutation inference (block length = 3 days).

## Results

### Hormone levels, sleep quality, and PVSV across the physiological menstrual cycle and OCP conditions

Progesterone levels were lower under OCP compared to the physiological menstrual cycle, with peak values reduced by approximately 97% (0·40 vs 15·50 ng/mL; p=0·016). Oestradiol, LH, and FSH did not differ between conditions (all p>0·34; **table 1**). 1-dPSQI global scores and all subcomponents did not differ between conditions (p=0·088 for the global score; all subcomponents p>0·24; **table 1**, **supplementary table 1**). Given that sleep medication was not used in any session, this subcomponent was excluded from further analyses. Total PVSV did not differ between conditions (p=0·88; **table 1**). Region-wise analysis showed that BG PVSV was lower under OCP compared to the physiological menstrual cycle (p=0·0051; **table 1**), whereas CSO PVSV did not differ between conditions (p=0·61). The hippocampus yielded zero PVSV across all sessions and was therefore excluded from further analyses.

**Table 1:** Descriptive statistics of hormones, PVSV, and subjective sleep quality across conditions. Values are presented as mean ± standard deviation [minimum, maximum]; variance ratio rows report only the F-test result, with corresponding SD values shown in the row above. One OCP day with an outlying 1-dPSQI value (|z|>3) was excluded from 1-dPSQI mean comparison. Hormone values are reported on the original scale; log-transformation was applied prior to statistical analysis. p-values from moving-block permutation tests (10,000 permutations, block length = 4 days, two-sided). CI, confidence interval; FSH, follicle- stimulating hormone; LH, luteinising hormone; 1-dPSQI, daily-adapted Pittsburgh Sleep Quality Index; PVSV, perivascular space volume. †p<0·10; *p<0·05; **p<0·01.

| Variable | Physiological menstrual cycle<br>(n = 30)<br>Mean (SD) [min–max] | Oral contraceptive pill<br>(n = 29)<br>Mean (SD) [min–max] | Mean difference<br>(95% CI) | p |
| --- | --- | --- | --- | --- |
| **Hormones** |  |  |  |  |
| Oestradiol<br>(pg/mL) | 82.84 ± 54.29<br>[21.80, 264.00] | 65.18 ± 73.34<br>[5.44, 246.00] | 17.66<br>(-36.61 to 67.02) | 0.56 |
| Progesterone<br>(ng/mL) | 5.14 ± 5.87<br>[0.02, 15.50] | 0.15 ± 0.12<br>[0.04, 0.40] | 4.99<br>(1.69 to 8.65) | **0.016\*** |
| LH<br>(mIU/mL) | 7.80 ± 7.80<br>[2.58, 45.27] | 6.04 ± 2.94<br>[2.10, 12.66] | 1.76<br>(-1.76 to 6.61) | 0.63 |
| FSH<br>(mIU/mL) | 5.82 ± 2.05<br>[3.08, 13.23] | 4.97 ± 1.91<br>[2.24, 9.03] | 0.86<br>(-0.70 to 2.40) | 0.34 |
| **Brain measure** |  |  |  |  |
| PVSV, total (mm<sup>3</sup>) | 1365.17 ± 99.64<br>[1163.00, 1580.00] | 1372.38 ± 126.63<br>[1138.00, 1647.00] | -7.21<br>(-86.52 to 64.35) | 0.88 |
| PVSV, CSO (mm<sup>3</sup>) | 908.37 ± 94.58<br>[732.00, 1115.00] | 930.38 ± 122.58<br>[713.00, 1220.00] | -22.01<br>(-97.74 to 45.74) | 0.61 |
| PVSV, BG (mm<sup>3</sup>) | 456.80 ± 17.05<br>[421.00, 481.00] | 442.00 ± 15.26<br>[419.00, 474.00] | 14.80<br>(8.03 to 21.32) | **0.0051\*\*** |
| **Sleep quality** |  |  |  |  |
| PSQI total score | 3.37 ± 2.19<br>[0.00, 9.00] | 2.38 ± 1.32<br>[0.00, 6.00] | 0.99<br>(-0.03 to 2.11) | 0.088† |
| **Day-to-day change (Δ)** |  |  |  |  |
| Δ PVSV, total<br>(mm<sup>3</sup>) | -0.72 ± 140.33<br>[-263.00, 278.00] | -7.19 ± 110.47<br>[-190.00, 239.00] | 6.46<br>(-28.34 to 42.84) | 0.70 |
| Δ 1-dPSQI total<br>score | -0.17 ± 2.75<br>[-5.00, 7.00] | 0.00 ± 1.54<br>[-2.00, 4.00] | -0.17<br>(-1.14 to 0.75) | 0.74 |
| Variance ratio,<br>Δ PVSV, total |  |  |  | 0.17 |
| Variance ratio,<br>Δ 1-dPSQI total<br>score |  |  |  | **0.019\*** |

### LH is associated with subjective sleep quality across the physiological menstrual cycle

To examine hormonal associations with sleep quality and PVSV during the physiological menstrual cycle, we fitted separate linear regression models on absolute daily values for each of the four hormones. Higher daily LH levels were associated with poorer subjective sleep quality (b=2·15, 95% CI 0·64 to 3·67, p_FDR_=0·028; **table 2**, **figure 2A**). However, the ovulatory day showed a high Cook’s distance in the 1-dPSQI–LH model, indicating disproportionated influence of this measurement on the model fit. We excluded this day to assess whether the finding was driven by a single observation and re-estimated the model. The association remained after exclusion (b=2·84, 95% CI 0·71 to 4·98, p=0·011). In contrast, oestradiol, progesterone, and FSH showed no association with 1-dPSQI scores (all p_FDR_>0·26; **supplementary figure 1A–C**). For PVSV, higher oestradiol levels were moderately associated with reduced PVSV (b=−61·70, 95% CI −118·38 to −5·02, p=0·034, p_FDR_=0·14; **table 2**, **figure 2B**), though this association did not survive FDR correction. Progesterone, LH, and FSH showed no association with PVSV (all p_FDR_>0·42; **supplementary figure 1D–F**).

**Figure 2:**
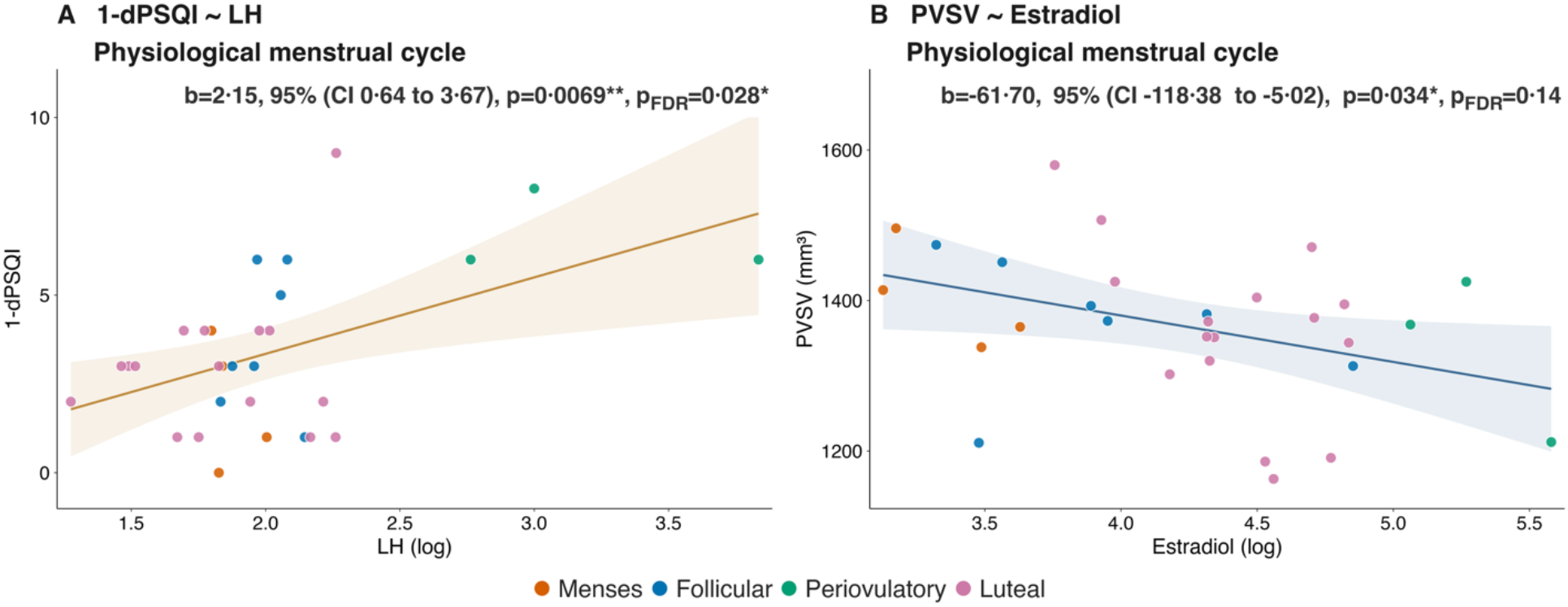
Associations between hormone levels, sleep quality and PVSV during the physiological menstrual cycle. **(A)** Linear regression of 1-dPSQI total score on log-transformed LH levels. (**B)** Linear regression of PVSV on log-transformed oestradiol levels. Data points are coloured by cycle phase. Regression lines with 95% confidence bands are shown. 95% CI, parametric confidence interval; p, uncorrected p-value; pFDR, Benjamini–Hochberg-corrected p-value; b, unstandardised regression coefficient. LH, luteinising hormone; 1-dPSQI, daily-adapted Pittsburgh Sleep Quality Index; PVSV, perivascular space volume. *p <0·05; **p <0·01.

**Table 2:** Regression models assessing hormone effects on subjective sleep quality and PVSV and hormonal moderation of the subjective sleep quality–PVSV association. Models 1 and 2 examine absolute hormone levels as predictors of 1-dPSQI total score and PVSV, respectively. Model 3 tests whether hormone levels moderate the association between 1-dPSQI total score and PVSV. All models were fitted using OLS regression (Durbin–Watson p>·14 for all models). FDR correction was applied separately per outcome group (Models 1 and 2) and across all interaction terms (Model 3). 95% CI derived using parametric confidence intervals. b, unstandardised regression coefficient; SE, standard error; 95% CI, 95% confidence interval; pFDR, Benjamini– Hochberg-corrected p-value. FSH, follicle-stimulating hormone; LH, luteinising hormone; 1-dPSQI, daily-adapted Pittsburgh Sleep Quality Index; PVSV, perivascular space volume. †p<0·10; *p<0·05; **p<0·01.

| Variable | *b* | SE | 95% CI | *p* | *p*<sub>FDR</sub> |
| --- | --- | --- | --- | --- | --- |
| **Model 1: 1-dPSQI ~ Hormone** |  |  |  |  |  |
| Oestradiol | 0.48 | 0.65 | -0.86 to 1.82 | 0.47 | 0.47 |
| Progesterone | -0.28 | 0.36 | -1.01 to 0.45 | 0.43 | 0.47 |
| LH | 2.15 | 0.74 | 0.64 to 3.67 | **0.0069\*\*** | **0.028\*** |
| FSH | 2.23 | 1.43 | -0.70 to 5.15 | 0.13 | 0.26 |
| **Model 2: PVSV ~ Hormone** |  |  |  |  |  |
| Oestradiol | -61.70 | 27.67 | -118.38 to -5.02 | **0.034\*** | 0.14 |
| Progesterone | -16.80 | 16.05 | -49.67 to 16.07 | 0.30 | 0.42 |
| LH | -5.08 | 38.35 | -83.64 to 73.48 | 0.90 | 0.90 |
| FSH | 68.36 | 66.51 | -67.88 to 204.60 | 0.31 | 0.42 |
| **Model 3: PVSV ~ 1-dPSQI × Hormone** |  |  |  |  |  |
| 1-dPSQI × Oestradiol | -1.20 | 11.94 | -25.74 to 23.35 | 0.92 | 0.92 |
| 1-dPSQI × Progesterone | 7.48 | 7.61 | -8.17 to 23.13 | 0.34 | 0.45 |
| 1-dPSQI × LH | -49.71 | 21.48 | -93.87 to -5.55 | **0.029\*** | 0.058† |
| 1-dPSQI × FSH | -92.98 | 32.68 | -160.15 to -25.81 | **0.0085\*\*** | **0.034\*** |

### Gonadotropin hormones moderate the sleep quality–PVSV association

To test whether reproductive hormones moderate the sleep quality–PVSV association rather than exerting direct main effects on PVSV, interaction models (PVSV ∼ 1-dPSQI × Hormone) were fitted at the absolute level. Among the gonadotropins, FSH moderated the sleep quality–PVSV association (b*=*−92·98, 95% CI −160·15 to −25·81, p_FDR_=0·034; **table 2**, **figure 3D**). A similar pattern was observed for LH but did not survive FDR correction (b=−49·71, 95% CI −93·87 to −5·55, p_FDR_=0·058; **figure 3E**). The negative interaction terms indicate that the sleep-associated increase in PVSV was attenuated at higher gonadotropin levels, with the 1-dPSQI–PVSV slope reversing during the periovulatory LH and FSH surge (**figure 3B – C**). Oestradiol and progesterone did not moderate the sleep quality–PVSV association (both p_FDR_>0·45; **table 2**).

**Figure 3:**
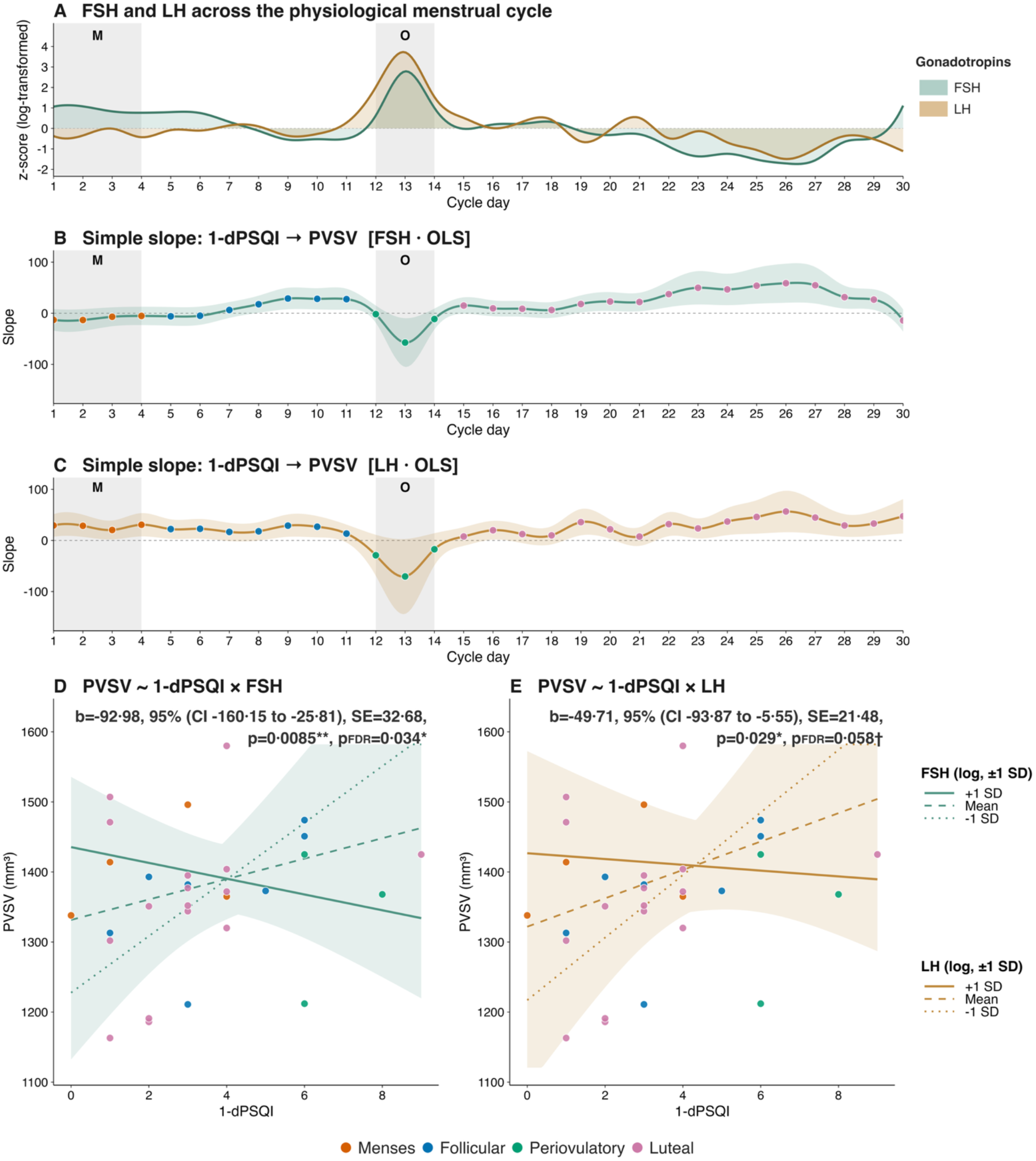
Gonadotropin-dependent modulation of the subjective sleep quality–PVSV association across the physiological menstrual cycle. (**A**) Z-standardised LH and FSH levels across the physiological menstrual cycle, shown as spline-smoothed trajectories connecting daily values. Grey-shaded areas indicate the menses (M) and the periovulatory window (O). (**B, C**) Simple slope of PVSV on 1-dPSQI derived from the interaction models PVSV ∼ 1-dPSQI × FSH (**B**) and PVSV ∼ 1-dPSQI × LH (**C**); lines and shaded confidence bands are spline-smoothed for visualisation, with data points representing the model-implied regression slope of 1-dPSQI on PVSV at the observed hormone level of that session. The shaded band represents the parametric 95% confidence interval of the conditional slope estimate. A positive slope indicates that poorer sleep quality is associated with larger PVSV on that day; a negative slope indicates an attenuation or reversal of this coupling. (**D, E**) Interaction plots for PVSV ∼ 1- dPSQI × FSH (**D**) and PVSV ∼ 1-dPSQI × LH (**E**). Regression lines are shown at ±1 SD and at the mean hormone level. Data points are coloured by cycle phase. The shaded band represents the parametric 95% confidence interval around the mean regression line. b, unstandardised interaction coefficient; 95% CI, parametric confidence interval; p, uncorrected p-value; pFDR, Benjamini– Hochberg-corrected p-value. FSH, follicle-stimulating hormone; LH, luteinising hormone; 1-dPSQI, daily-adapted Pittsburgh Sleep Quality Index; PVSV, perivascular space volume; OLS, ordinary least squares. †p <0·10; *p <0·05; \**\**p <0·01.

### Positive association between day-to-day changes in sleep quality and PVSV is specific to the physiological menstrual cycle

To capture acute, synchronous changes between sleep quality and PVSV, we computed Spearman rank correlations on day-to-day differences (Δ1-dPSQI, ΔPVSV) within each condition. During the physiological menstrual cycle, day-to-day decreases in sleep quality were associated with synchronous increases in PVSV (ρ=0·55, 95% CI 0·30 to 0·67, p_perm_=0·0095; **figure 4C**). This association was primarily driven by the subcomponent ‘subjective sleep quality’ of the 1-dPSQI (**supplementary figure 3**). This association was absent under OCP (ρ=−0·17, 95% CI −0·58 to 0·46, p_perm_=0·44; **figure 4D**). The association was different between conditions (Δρ=0·72, p=0·040), suggesting that PVSV expansion following nights of poorer sleep quality is specific to the physiological menstrual cycle.

**Figure 4:**
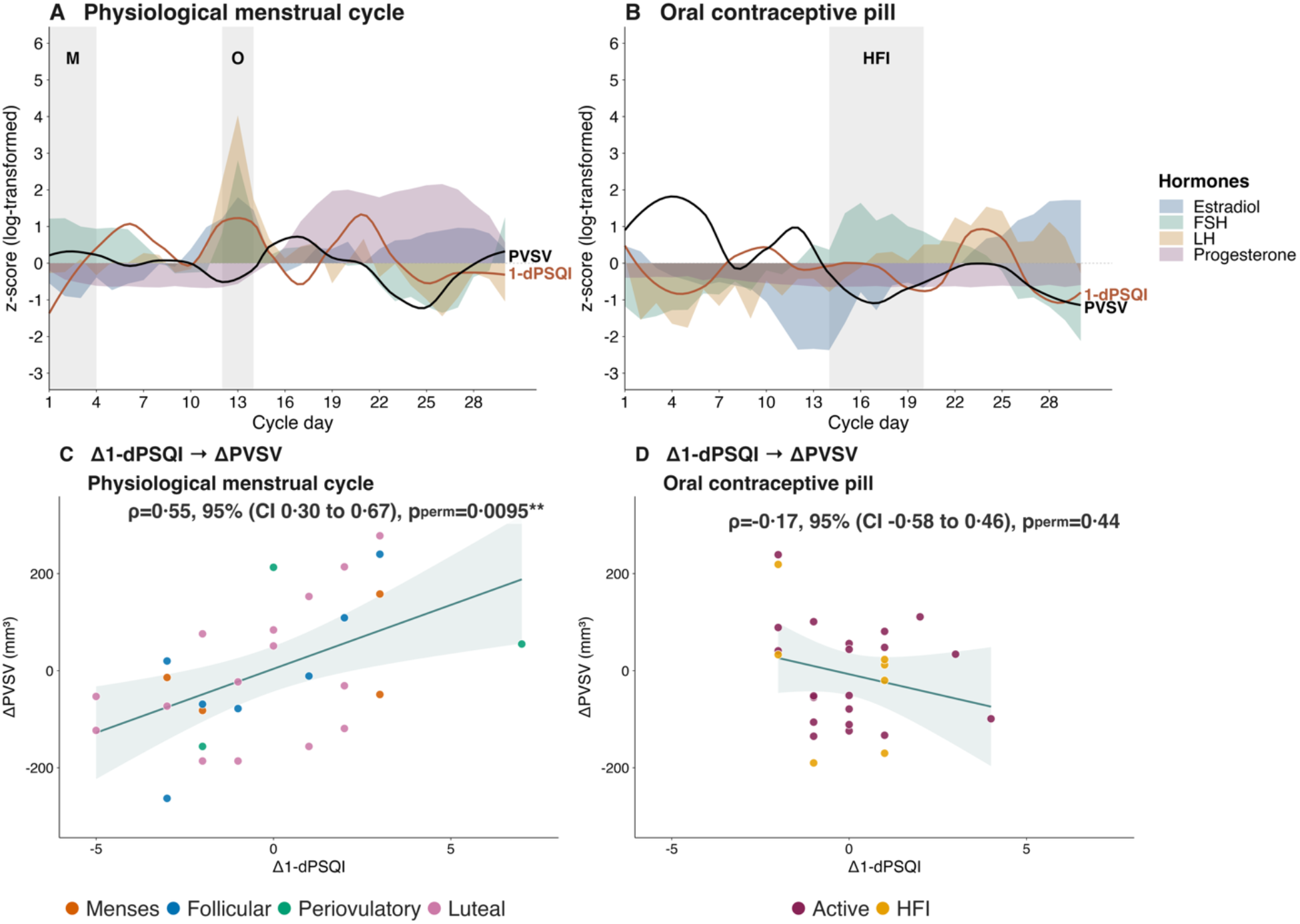
Daily dynamics of subjective sleep quality and PVSV across the physiological menstrual cycle and under oral contraceptive pill use. **(A, B)** Z-standardised time series of PVSV and 1-dPSQI total score across the 30-session measurement period for the physiological menstrual cycle (**A**) and the OCP condition (**B**). PVSV and 1-dPSQI trajectories are displayed as LOESS-smoothed curves (span = 0.35). Hormone trajectories are displayed as shaded ribbons (log-transformed, z-scored). Grey shaded areas indicate the menses (M) and periovulatory window (O) in the physiological menstrual cycle and the hormone-free interval (HFI) in the OCP condition. Dashed horizontal line indicates z = 0. **(C, D)** Day-to-day changes in 1-dPSQI total score (Δ 1-dPSQI) and PVSV (Δ PVSV) in the physiological menstrual cycle (**C**) and the OCP condition (**D**). Data points are coloured by cycle phase. Shaded band represents the 95% confidence interval. ρ, Spearman rank correlation coefficient; CI, bootstrap-based confidence interval (10,000 iterations); p, permutation-based p-value (10,000 permutations, two-sided). FSH, follicle-stimulating hormone; LH, luteinising hormone; OCP, oral contraceptive pill; 1-dPSQI, daily-adapted Pittsburgh Sleep Quality Index; PVSV, perivascular space volume. **p <0·01.

Given the lower day-to-day 1-dPSQI variability under OCP than during the physiological menstrual cycle (variance ratio=3·18, p_perm_=0·019; **table 1**), we conducted a sensitivity analysis to assess whether this difference could be attributable to range restriction of 1-dPSQI during OCP use. Physiological menstrual cycle data were resampled within the Δ1-dPSQI range observed during OCP use (5000 bootstrap replicates). The resulting coefficients (median ρ=0·57, 95% percentile interval 0·23–0·80) aligned with the unrestricted physiological menstrual cycle estimate, indicating that the absence of the Δ1-dPSQI–ΔPVSV association under OCP reflects a difference between conditions rather than an artefact of reduced sleep quality variability.

Lagged cross-correlation analyses confirmed the temporal specificity of the Δ1-dPSQI–ΔPVSV association, with no associations observed at lags +1 to +3 days (all p_FDR_>0·44), indicating that the association did not extend to delayed effects of sleep quality on PVSV (**supplementary table 2**).

Regional PVSV analyses revealed that this association was primarily driven by CSO-PVSV (r_s_=0·64, 95% CI 0·45 to 0·76, p_perm_=0·0017), whereas BG-PVSV showed no significant day-to-day association with sleep quality (ρ=−0·25, 95% CI −0·62 to 0·14, p_perm_=0·29). Partial correlations confirmed that the sleep quality– PVSV association remained after controlling for BG-PVSV (r_partial_=0·74, 95% CI 0·53 to 0·87, p_perm_=0·0002) but was reversed after controlling for CSO-PVSV (r_partial_=−0·53, 95% CI −0·76 to −0·15, p_perm_=0·021), indicating that CSO-PVSV accounts for the observed sleep quality–PVSV association (**supplementary figure 2**).

## Discussion

Using a single-subject dense-sampling design, we examined the influence of sustained endocrine effects and day-to-day hormonal fluctuations on subjective sleep quality, PVSV, and their association. Our main findings demonstrated that *i)* during the physiological menstrual cycle, higher LH levels were associated with poorer sleep quality. For PVSV, a negative association with oestradiol emerged, although it did not survive FDR correction; *ii)* The association between sleep quality and PVSV during the physiological menstrual cycle was further moderated by gonadotropins. In particular, higher FSH levels attenuated the association between poorer sleep quality and increased PVSV, with a comparable trend in LH. Given that both FSH and LH rise during the periovulatory window, this moderating effect may reflect an interplay between gonadotropins, sleep quality, and CSF flow into the brain parenchyma, indicating an ovulation- specific attenuation of the sleep quality–PVSV association; *iii)* Sleep quality showed greater day-to-day variability within the physiological menstrual cycle than during OCP use. Moreover, greater declines in sleep quality were associated with greater increases in PVSV during the physiological menstrual cycle, an association absent in the OCP condition.

Within the physiological menstrual cycle, higher LH levels were associated with poorer sleep quality. This is supported by former studies reporting a positive association between LH levels and nocturnal awakenings.^3^ LH levels start rising within the follicular phase, characteristically surging during the periovulatory window. This surge is driven by kisspeptin-induced gonadotropin-releasing hormone secretion, while noradrenergic release from the LC is thought to further modulate both the timing and magnitude of gonadotropin release.^19^ Enhanced noradrenergic LC activity, particularly reflected in high-amplitude noradrenergic surges, was shown to promote microarousals and NREM sleep fragmentation,^20^ suggesting that the association between elevated LH levels and poorer sleep quality may reflect noradrenergically mediated sleep alterations during the periovulatory window. Additionally, we observed a pronounced decrease in PVSV within the periovulatory window, as reflected by a moderating effect of gonadotropins on the sleep quality–PVSV association. Enhanced noradrenergic activity accompanying the LH surge may further promote parenchymal resistance to CSF bulk flow,^21^ thereby reducing PVSV, and ultimately altering CSF inflow. Interestingly, a reduction in CSF volume at the timepoint of ovulation has been reported previously.^22^ While this study assessed structural CSF compartments rather than CSF inflow, it may align with our assumption that CSF inflow is altered within the periovulatory window.

We further observed a moderate association between lower PVSV and higher oestradiol levels. Oestradiol enhances cerebral perfusion by potentiating endothelial NO signaling, which relaxes vascular smooth muscle and thereby reduces cerebral vascular tone.^23^ A positive association between cerebral perfusion and oestradiol in naturally cycling women is reported.^24^ Enhanced perfusion-induced vasodilation has been shown to reduce glymphatic CSF inflow in a murine model, most likely due to reduced PVS luminal area.^11^

Given the characteristic rise in oestradiol during the periovulatory window, this vasodilatory mechanism may act alongside noradrenergic modulation, contributing to the observed reduction in PVSV.

In the physiological menstrual cycle, poorer sleep quality was associated with higher PVSV in the CSO. Previous studies found that one night of poor sleep is associated with insufficient dissipation of sleep pressure, resulting in elevated residual sleep pressure the following morning.^25^ Accordingly, elevated sleep pressure following sleep deprivation was linked with increased vasomotion.^26^ Our data demonstrated that even modest reductions in sleep quality may increase residual sleep pressure, thereby promoting vasomotion and enhancing CSF inflow, as reflected by the observed increase in PVSV. However, whereas mean sleep quality did not differ between the physiological menstrual cycle and OCP condition, its day-to- day variability was significantly lower under OCP. Interestingly, OCP use is associated with suppressed fluctuations in affective symptoms, most likely due to attenuated hypothalamic–pituitary–adrenal axis reactivity.^27^ While the underlying mechanism is not fully understood, it may involve reduced central corticotropin-releasing hormone release, a neuropeptide that promotes LC activation and enhances arousal-related sleep alterations.^28^ This reduced day-to-day variability in sleep quality may diminish the incidence of insufficient sleep pressure dissipation, thereby attenuating sleep-pressure related vasomotion and the associated increase in PVSV.

The intraindividual design provides implicit control for stable between-subject confounds, including interindividual response tendencies and vascular and physiological characteristics known to influence PVS morphology,^10^ while enabling examination of sleep quality–PVSV coupling across hormonal milieus under reduced confounding. Moreover, MRI measurements were performed at a constant time in the morning, thereby controlling for the time-of-day effects on PVSV.^29^ Despite the advantages of dense sampling, several limitations should be mentioned. The single-subject design limits generalisability and requires replication across different datasets and diverse hormonal milieus. Our PVS segmentation relied solely on T1-weighted images, while using T2/FLAIR imaging and the Frangi Filter method could improve PVS delineation. Finally, PVSV provides a structural proxy of CSF influx, but does not directly assess solute clearance or exchange. Future studies could incorporate innovative non-invasive approaches such as parenchymal impedance spectroscopy,^7^ which infers extracellular fluid volume from tissue conductivity, or CSF-STREAM, which enables measurement of CSF mobility.^30^ Finally, we used self-reported sleep quality, whereas polysomnography or actigraphy would provide an objective characterisation of sleep architecture.

Our findings reveal that the periovulatory window is a sensitive period within the physiological menstrual cycle in which the sleep quality–PVSV interaction is altered, likely driven by noradrenergic modulation.

We further showed that cyclical hormonal changes are associated with day-to-day variability in sleep quality, whereas these fluctuations are attenuated under hormonal suppression. We also showed that PVSV increase was associated with poor sleep quality, possibly reflecting elevated sleep pressure-related CSF influx. Together, we found that the menstrual cycle may constitute a physiologically active state relevant to sleep health and brain physiology, which is altered by hormonal suppression. Given that glymphatic function is a promising target in brain diseases, future studies should explore in vivo sleep-related brain clearance through the lens of hormonal modulation in women.

## Data sharing

The study leveraged the publicly available ‘28andMe’ dataset, which is available on OpenNeuro (https://openneuro.org/datasets/ds002674/versions/1.0.6). Raw 1-dPSQI values were obtained from the dataset authors upon request.

## Code availability

Code for the neuroimaging preprocessing, PVSV calculation, and statistical analysis is available on GitHub (https://github.com/d-apter/sleep_pvs_cycle).

## Acknowledgments

We would like to thank Laura Pritchet for collecting the ‘28andMe’ dataset, making it publicly available, and providing additional data upon request.

