## Supplementary material for "Hormonal Modulation of Subjective Sleep Quality and Perivascular Space Volume Across the Menstrual Cycle: An Observational Dense-Sampling Study"

### Supplementary Tables & Figures


#### Supplementary Tables……………………………………………………………...……2

Supplementary Table 1: Descriptive statistics of 1-dPSQI subcomponent scores across conditions…………..…...…2

Supplementary Table 2: Lagged cross-correlations between day-to-day changes in 1-dPSQI global score and PVSV……………………………………………………………………………………………….……….3

#### Supplementary Figures…………...…………………...…………………………...……4

Supplementary Figure 1: Non-significant associations between hormone and subjective sleep quality and PVSV in the physiological menstrual cycle……………………………………………………………………...…4

Supplementary Figure 2: Regional specificity of the day-to-day subjective sleep quality–PVSV association in the physiological menstrual cycle……………………………………………………………………..…..5

Supplementary Figure 3: Spearman's rank correlations between PVSV and 1-dPSQI subcomponent scores across conditions……………………………………………………………………………………………6

Supplementary Figure 4: Spearman rank correlations between hormone levels and 1-dPSQI subcomponent scores in the physiological menstrual cycle……………………………………………………………………..7

#### Supplementary Tables

| 1-dPSQI subcomponent | Physiological menstrual cycle (n = 30) Mean (SD) [min–max] | Oral contraceptive pill (n = 29) Mean (SD) [min–max] | Mean difference (95% CI) | p |
|---|---|---|---|---|
| Subjective sleep quality | 0·77 ± 0·73 [0·00, 2·00] | 0·63 ± 0·85 [0·00, 3·00] | 0·13 (-0·30 to 0·57) | 0·67 |
| Sleep latency | 0·53 ± 0·78 [0·00, 2·00] | 0·37 ± 0·56 [0·00, 2·00] | 0·17 (-0·27 to 0·57) | 0·51 |
| Sleep duration | 0·27 ± 0·45 [0·00, 1·00] | 0·17 ± 0·38 [0·00, 1·00] | 0·10 (-0·13 to 0·33) | 0·58 |
| Habitual sleep efficiency | 0·20 ± 0·48 [0·00, 2·00] | 0·03 ± 0·19 [0·00, 1·00] | 0·17 (-0·00 to 0·37) | 0·24 |
| Sleep disturbances | 0·70 ± 0·47 [0·00, 1·00] | 0·53 ± 0·51 [0·00, 1·00] | 0·17 (-0·13 to 0·47) | 0·43 |
| Daytime dysfunction | 0·70 ± 0·75 [0·00, 3·00] | 0·83 ± 0·87 [0·00, 3·00] | -0·13 (-0·40 to 0·13) | 0·60 |

**Supplementary Table 1: Descriptive statistics of 1-dPSQI subcomponent scores across conditions.** Values are presented as mean ± standard deviation [minimum, maximum]. One OCP day with an outlying 1-dPSQI value (|z|>3) was excluded. Scores range from 0 to 3 per subcomponent, with higher scores indicating poorer sleep quality. Subcomponent sleep medication use was excluded from analysis due to zero variance across all sessions. 95% CI derived from bootstrap resampling (10,000 iterations, length = 3 days). p-value from a moving-block permutation test (10,000 permutations, block length = 4 days, two-tailed) for the difference in means between conditions. 1-dPSQI, daily-adapted Pittsburgh Sleep Quality Index.

| | Physiological menstrual cycle | | | Oral contraceptive pill | | |
|---|---|---|---|---|---|---|
| Lag (days) | n | ρ (95% CI) | $p_{FDR}$ | n | ρ (95% CI) | $p_{FDR}$ |
| +1 | 28 | -0·32 (-0·67 to 0·11) | 0·44 | 26 | 0·04 (-0·45 to 0·44) | 0·96 |
| +2 | 27 | 0·26 (-0·08 to 0·61) | 0·44 | 25 | -0·01 (-0·51 to 0·42) | 0·96 |
| +3 | 26 | -0·13 (-0·53 to 0·24) | 0·59 | 24 | -0·06 (-0·41 to 0·36) | 0·96 |

**Supplementary Table 2: Lagged cross-correlations between day-to-day changes in 1-dPSQI global score and PVSV.** Positive lags indicate that Δ1-dPSQI precedes ΔPVSV by the respective number of days. 95% CI derived from block-bootstrap resampling (10,000 iterations, block length = 3 days). $p_{FDR}$ values reflect Benjamini–Hochberg-corrected p-values from block-permutation-based inference (10,000 permutations, block length = 3 days, two-sided), corrected within each condition's three-lag family (lags +1 to +3). OCP, oral contraceptive pill; 1-dPSQI, daily adapted Pittsburgh Sleep Quality Index; PVSV, perivascular space volume.

#### Supplementary Figures

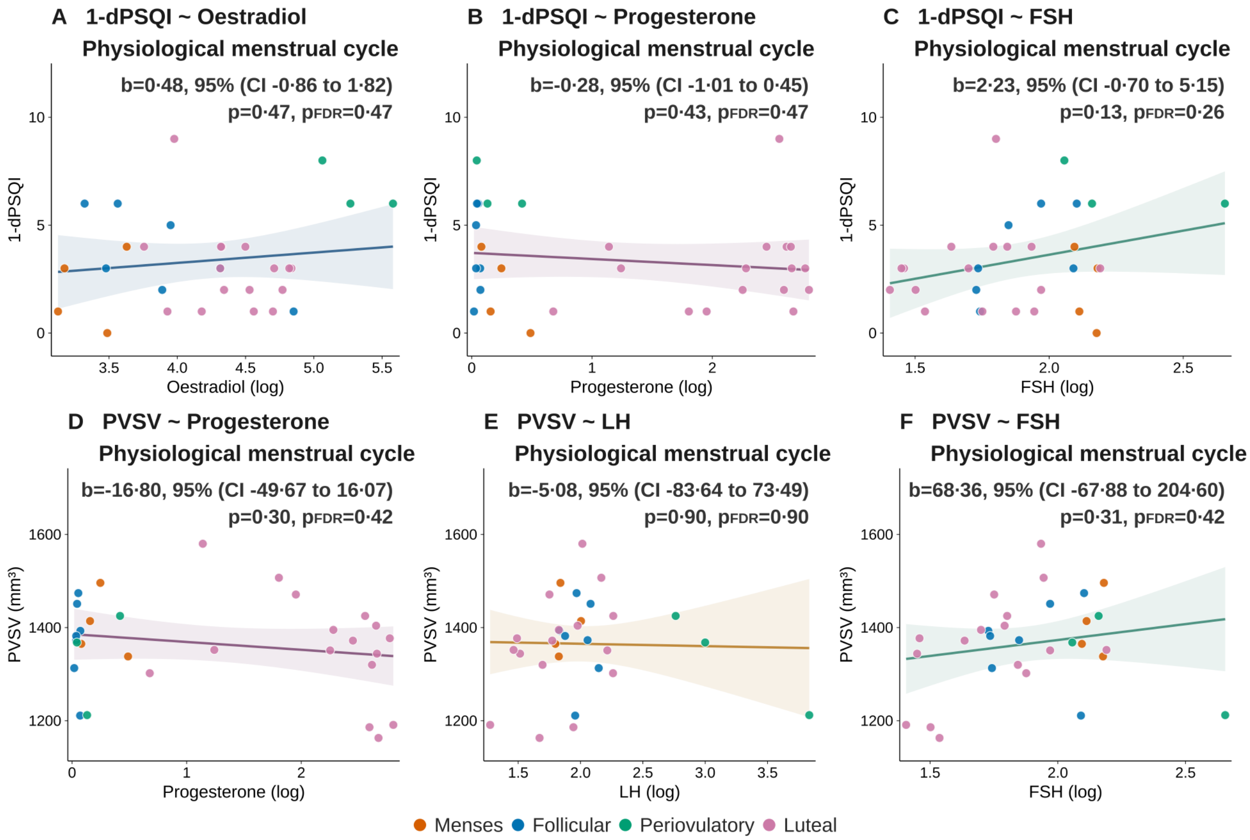


**Supplementary Figure 1: Non-significant associations between hormone and subjective sleep quality and PVSV during the physiological menstrual cycle.** (**A–C**) Linear regression of 1-dPSQI total score on log-transformed oestradiol (**A**), progesterone (**B**), and FSH (**C**) levels. (**D–F**) Linear regression of PVSV on log-transformed progesterone (**D**), LH (**E**), and FSH (**F**) levels. Data points are coloured by cycle phase. Regression lines with parametric 95% confidence bands are shown. None of the displayed associations survived correction for multiple comparisons. b, unstandardised regression coefficient; 95% CI, parametric confidence interval; p, uncorrected p-value; $p_{FDR}$, Benjamini–Hochberg-corrected p-value. FSH, follicle-stimulating hormone; LH, luteinising hormone; 1-dPSQI, daily-adapted Pittsburgh Sleep Quality Index; PVSV, perivascular space volume.

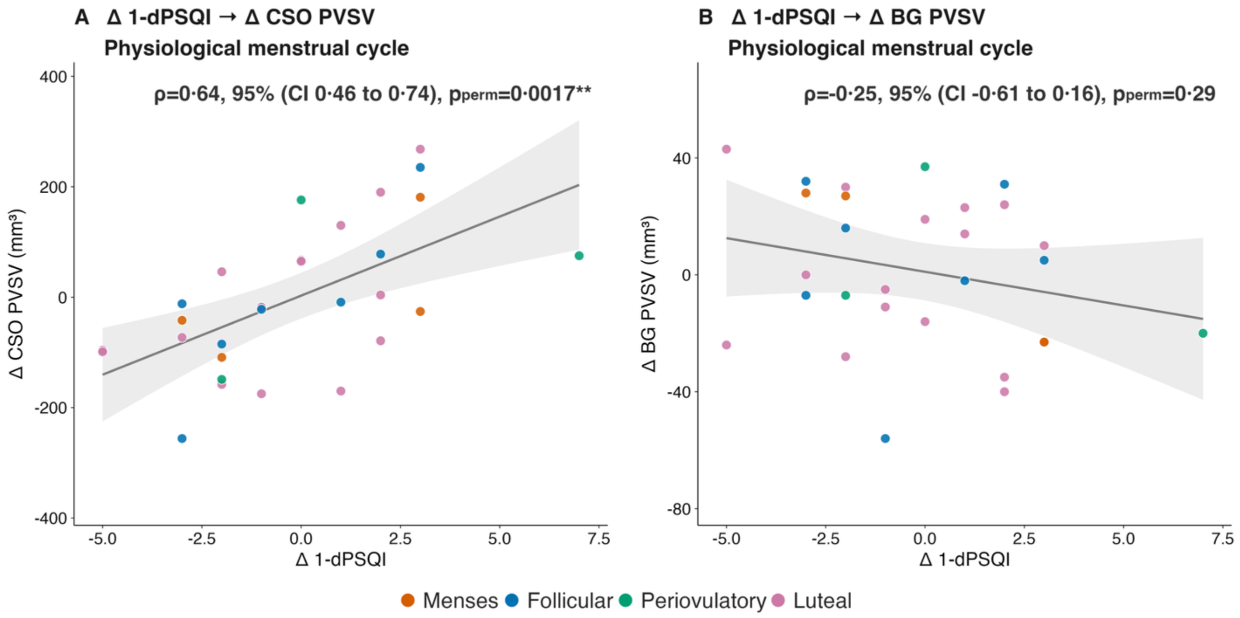


**Supplementary Figure 2: Regional specificity of the day-to-day subjective sleep quality–PVSV association during the physiological menstrual cycle. (A, B)** Spearman rank correlations between day-to-day changes in 1-dPSQI global score (ΔPSQI) and PVSV in the centrum semiovale (ΔCSO PVS; **A**) and basal ganglia (ΔBG PVS; **B**). Data points are coloured by cycle phase. The centrum semiovale showed a positive coupling with sleep quality, whereas no significant association was observed for the basal ganglia. Shaded band represents the 95% confidence interval. ρ, Spearman rank correlation coefficient; CI, bootstrap-based confidence interval (10,000 iterations); p, permutation-based p-value (10,000 permutations, two-sided). 1-dPSQI, Pittsburgh Sleep Quality Index; PVSV, perivascular space volume; CSO, centrum semiovale; BG, basal ganglia. **p<0·01.

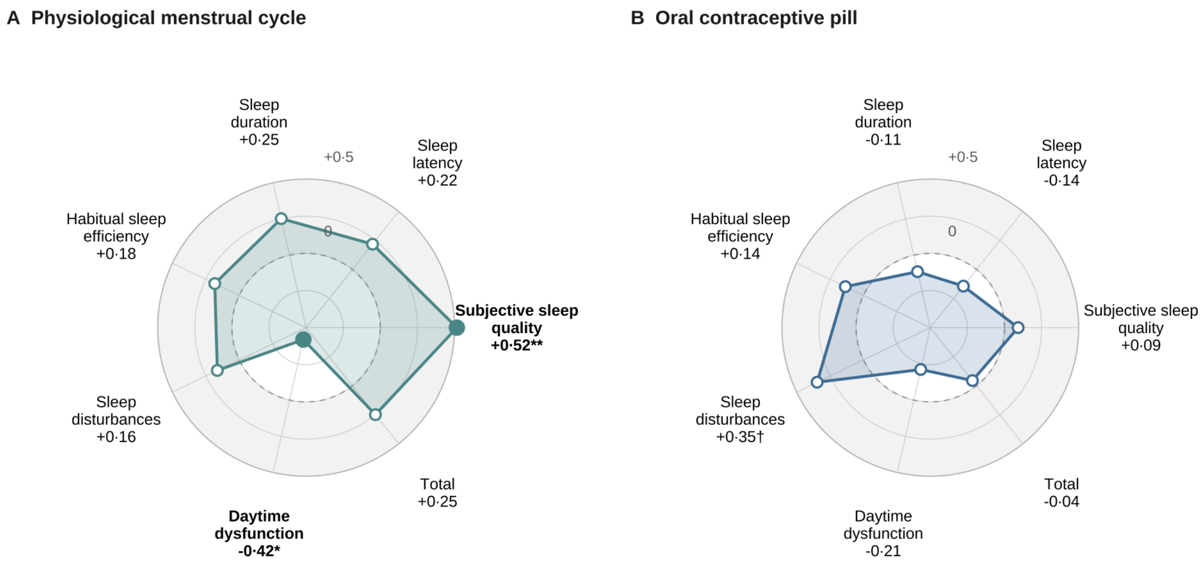


**Supplementary Figure 3: Spearman's rank correlations between PVSV and 1-dPSQI subcomponent scores across conditions. (A, B)** Radar plots displaying Spearman correlation coefficients between PVSV and six 1-dPSQI subcomponents and global score for the physiological menstrual cycle (**A**) and the OCP condition (**B**). Subcomponent sleep medication use was excluded from analysis due to zero variance across all sessions. Grey shading indicates positive Spearman correlation coefficients (ρ>0); the white inner zone indicates negative coefficients (ρ<0), separated by the dashed zero-reference circle. Filled markers indicate statistically significant correlations (p<0·05); open markers indicate non-significant correlations. These analyses are exploratory and were not corrected for multiple comparisons. OCP, oral contraceptive pill; 1-dPSQI, daily-adapted Pittsburgh Sleep Quality Index; PVSV, perivascular space volume. †p <0·10; ; *p<0·05; **p<0·01.

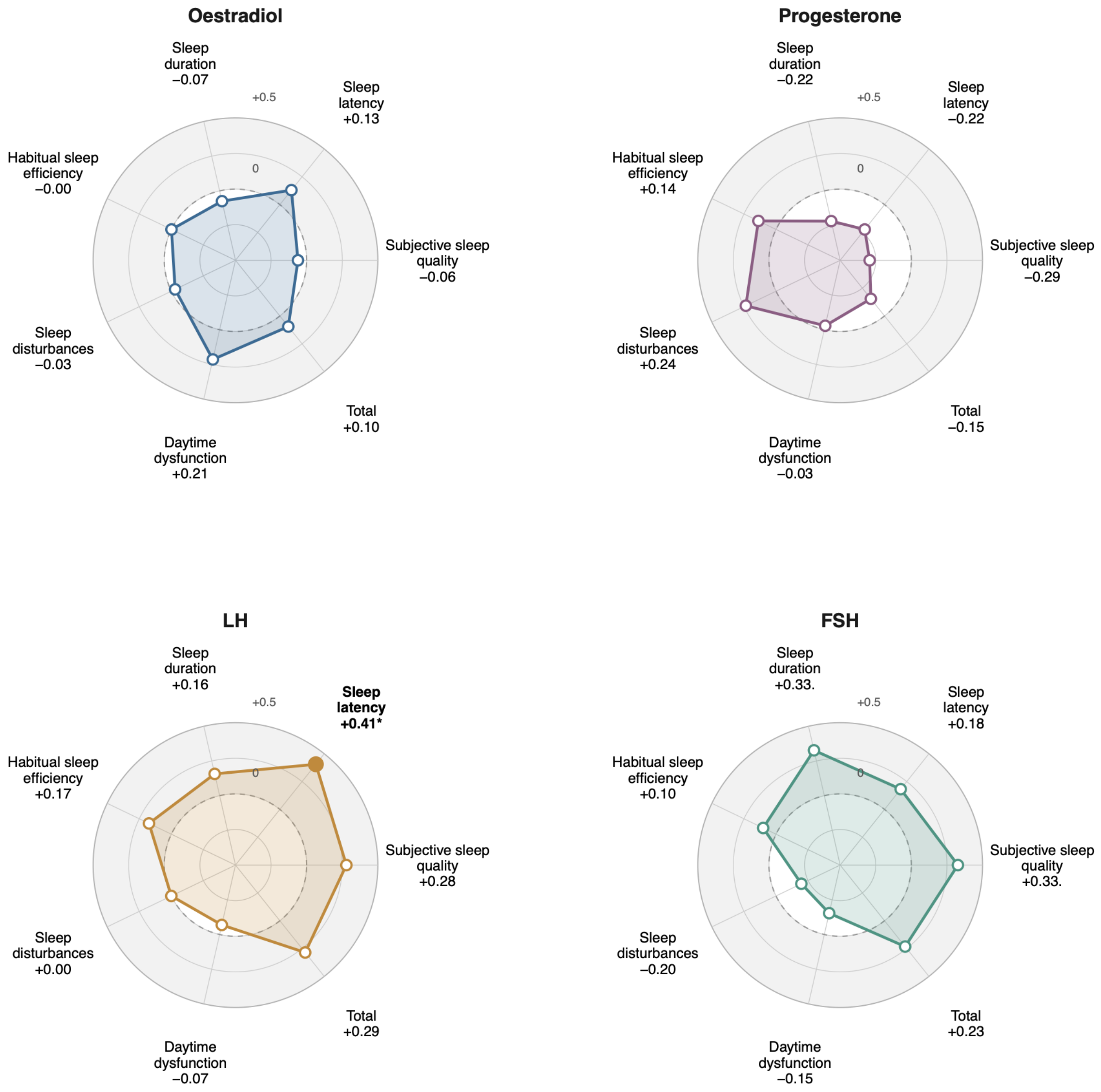


**Supplementary Figure 4: Spearman rank correlations between hormone levels and 1-dPSQI subcomponent scores in the physiological menstrual cycle.** Radar plots displaying Spearman correlation coefficients between each of the four hormones (oestradiol, progesterone, LH, FSH) and six 1-dPSQI subcomponents and global score. Subcomponent sleep medication use was excluded from analysis due to zero variance across all sessions. Each axis represents one 1-dPSQI subcomponent. Grey shading indicates positive Spearman correlation coefficients (ρ>0); the white inner zone indicates negative coefficients (ρ<0), separated by the dashed zero-reference circle. Filled markers indicate statistically significant correlations (p<0·05); open markers indicate non-significant correlations. These analyses are exploratory and were not corrected for multiple comparisons. FSH, follicle-stimulating hormone; LH, luteinizing hormone; OCP, oral contraceptive pill; 1-dPSQI, daily-adapted Pittsburgh Sleep Quality Index. †p<0·10; *p<0·05.
